# Predicting the genetic loci of past evolution

**DOI:** 10.1101/205153

**Authors:** Virginie Courtier-Orgogozo, Arnaud Martin

## Abstract

Repetitions in the mutations found to be responsible for independent evolution of similar phenotypes in various taxa have led some biologists to propose that for certain evolutionary changes the causal mutations are predictable. We examine here the nature of the predictions that have been made and their associated arguments. Predictions about the loci of past evolution are retrodictions, i.e. inferences about events that occurred in the past. They are not based on elaborate models and they derive mainly from the observation of repeated cases of genetic evolution. Predictions at the nucleotide level or at the gene level have a higher inference gain than those for broader categories of genetic changes such as cis-regulatory mutations.

## Introduction

Evolution reveals itself by the changes in observable characteristics of biological populations over successive generations. Here we focus on the DNA mutations underlying phenotypic changes that have occurred during natural evolution of populations or species, as well as through domestication and experimental evolution. The search for the mutations responsible for evolutionary changes started in the 70es, with iconic case studies such as the ABO blood group gene (F.I. YAMAMOTO et al. 1990) or hemoglobin for drepanocytosis and malaria resistance (V.M. INGRAM 1957). As instances of genes causing phenotypic changes between populations and species started to accumulate, certain researchers noticed that the mutations causing evolution did not appear to be randomly distributed across the genomes. Intriguing cases of repeated evolution at the genetic level were reported, with recurrent genetic changes involved in the evolution of similar phenotypes in distant taxa (A.H. PATERSo*n et a*l. 1908; N. GOMPEL and B. PRUD’HOMME 2009; A. KOPP 2009; P.A. CHRISTIN *et al.* 2010; A.E. LOBKOVSKY and E.V. KOONIN 2012; G.L. CONTE *et al.* 2012; A. MARTIN and V. ORGOGOZO 2013; D.L. STERN 2013; T. LENSER and G. THEISSEN 2013). Furthermore, certain types of phenotypic changes seemed to be preferentially associated with certain broad categories of mutations (S.B. CARROLL 2000; E.H. DAVIDSON 2006; G.A. WRAY 2007; B. PRUD'HOMME; N. GOMPEL and S.B. CARROLL 2007; S.B. CARROLL 2008; D.L. STERN and V. ORGOGOZO 2008; V.J. LYNCH and G. WAGNER 2008; G. WAGNER and V.J. LYNCH 2008; M.A. STREISFELD and M.D. RAUSHER 2011; D.L. STERN 2011). For example, morphological evolution in animals was suggested to preferentially involve cis-regulatory mutations rather than coding changes. Please note that “preferentially” refers here to the final consequence of selection and of other population genetics processes and does not necessarily mean that mutations occur in a non-random fashion. Genetic variations are usually thought to occur randomly throughout the genome, some of them being gradually eliminated while others are maintained, thus allowing enrichments in certain types of mutations when one looks at the result of selection and of other population genetics processes over multiple generations.

Observation of consistent patterns in the genetic loci of evolution has had two main consequences on evolutionary biology research. First, it prompted the elaboration of various explanatory hypotheses. Second, it led some biologists to propose that the genetic loci of evolution are partly predictable, in the sense that for a given phenotypic change that occurred in the past the underlying mutations can be guessed with reasonable confidence What is predicted here is the genetic causes of evolutionary events that occurred in the past (V. ORGOGOZO 2015; V. ORGOGOZO, B. MORIZOT and A. MARTIN 2015), and not the mutations that will occur in the future (for such cases see LOBKOVSKY and KOONIN 2012; R.E. LENSKI 2017; M. LÄSSING, V. MUSTONEN AND A.M. WALCZAK 2017).

In this paper we examine the predictions regarding the genetic loci of past evolution: what kinds of predictions are they? What are they based on? How accurate are they?

## Nature of the Predictions

Published predictions about the genetic loci of evolution do not arise from complex mechanistic models. They simply derive from the observation of repeated cases of genetic evolution and the identification of the context and the hypothetical causes that lead to repetitions. For example, the loss of larval trichomes in *Drosophila sechellia* was found to be caused by multiple mutations in five distinct enhancers of the *svb* gene, with each enhancer regulating trichome development in a specific body part (N. FRANKEL *et al.* 2010). Such recurrent evolution at the same locus, together with the special “hub” position of *svb* in the gene network for trichome development, suggested that *svb* was a hotspot gene for trichome evolution in flies (A.P. MCGREGOR *et al.* 2007; STERN and ORGOGOZO 2008; STERN 2011). The independent loss of trichomes in *Drosophila ezoana* was thus inferred to involve cis-regulatory mutations in *svb*, as in *D. sechellia*, and indeed this was found to be true (N. FRANKEL, S. WANG and D.L. STERN 2012). Agricultural pests and mosquitoes have repeatedly evolved resistance to pyrethroid insecticides such as DDT via coding mutations in the *para voltage-gated sodium channel* gene (*syn. para, vgsC*) (D.M. SODERLUND 2008). Weston et al. predicted that pyrethroids may also affect non-pest organisms that populate sprayed areas (D.P. Weston *et al.* 2013). Not only they found that pyrethroid resistance had evolved in populations of freshwater crustaceans exposed to agricultural run-off, but they also uncovered typical mutations in *para* gene, mirroring the evolutionary mechanisms previously observed in the targeted pests. Thus, a specific evolutionary pressure left a predictable genetic signature in the environment that can now be detected. In principle, comparable predictions could be done in other situations where human activity has chemically modified the environment.

Predictions rely on the assumption that the set of already known loci of evolution on which predictions are based are identified via an unbiased approach. Linkage mapping studies and association studies screen the entire genome sequence and are thus supposed to be unbiased in their detection of the genetic loci. However, one should keep in mind that once a genomic region is narrowed down to a few candidate genes through linkage mapping or association mapping, knowledge from past studies might favor for validation tests the candidate genes that are already known to be involved in a similar phenotypic change. As a result, even genome-wide mapping studies carry some bias towards already known genes.

Predictions are usually formulated along the following lines: “For a given phenotypic change, it is predicted that the causal mutations are such and such”. Formulations can also be relatively more complex. For evolution of red flowers in *Penstemon barbatus*, Wessinger and Rausher not only predicted the causal genes (*F3'5'H* or *F3'H*) and the fact that the mutations should create loss-of-function alleles but also provided gene-specific details about the expected mutations: “when it involves elimination of F3'5'H activity, functional inactivation or deletion of this gene tends to occur; however, when it involves elimination of F3'H activity, tissue-specific regulatory substitutions occur and the gene is not functionally inactivated” (C.A. WESSINGER and M.D. RAUSHER 2014).

## Predicting the genetic loci of evolution is a retrodiction

Predictions are usually inferences about the future, based on current knowledge about the past. When the causal temporality is reversed, some authors prefer to use the term retrodiction. Providing a *post hoc* explanation for an already known fact about the past is thus considered a retrodiction. For example, Darwin retrodicted why species similar to those found on oceanic islands are usually found on the nearest mainland (C. DARWIN 1859). More generally, retrodicting can be defined as making an inference about an event that occurred in the past (A. LOVE, personal communication; J. FETZER 2017). If this past event is already known, then retrodiction is the act of providing an explanation for it. Predictions are not always explanatory: they do not necessarily rely on a model or on causal explanations. For example, a prehistorical astronomer without a heliocentric theory who accumulates observations such as "the sun always rises above that hill" would predict that the sun will rise above the hill again, simply by noticing the repetitions. In our case, published predictions about the genetic loci of evolution do not arise from complex mechanistic models, they instead derive from the observation of repeated cases of genetic evolution. In this sense, evolutionary geneticists resemble prehistorical astronomers without a heliocentric theory. Using repetition among the known loci of evolution to make predictions about the past are thus retrodictions. Moreover, predictions about past genetic loci concern a genetic difference that exists today between two living taxa (a property of the present state). Predictions about the genetic loci of past evolution are thus retrodictions about the past and the present.

## Predictions at various genetic levels

Predictions can be made at the gene level, as for *svb* or the *para* sodium channel gene, but also at higher and lower genetic levels: at the level of a nucleotide, of part of a coding region, of a specific enhancer, of a group of genes, and also for broader categories of genetic changes (Table 1). Some predictions carry more information than others. Predicting the impact of a projectile within 1 m is better than within 1 km. In a first approximation, the *inference gain* can be estimated by the inverse of the probability of the predicted outcome according to the null model. In most cases, the null model considers that mutations can occur with equal probability at all nucleotide positions within a genome (ORGOGOZO 2015), or with equal probability in all genes of a genome. The prediction that trichome pattern evolution in *Drosophila* is likely to involve the *svb* gene (MCGREGOR *et al.* 2007) has an inference gain of approximately 17,500 (*svb* is one gene out of the total of 17,737 genes present in the genome of *D. melanogaster* (FlBase FB2017_04)). In contrast, the prediction that morphological evolution is likely to involve signaling ligand encoding genes (A. MARTIN and V. ORGOGOZO 2017) has a lower inference gain, estimated to be about 180 in the stickleback fish, as the proportion of genes associated with gene ontology (GO) terms “extracellular region” and “receptor binding” in the stickleback *G. aculeatus* (BROADS1) is 115/20,787 (MARTIN and ORGOGOZO 2017). Predictions at the level of the gene or at the nucleotide level have a higher inference gain than predictions about broader classes of mutations.

**Table 1.** **Repetitions can be detected and predicted at various genetic levels, from specific nucleotides to general classes of mutations.**

| Genetic Level | Example of prediction or of repeated evolution |
| --- | --- |
| Nucleotide | Resistance to cyclodiene insecticides was successfully predicted to be associated with amino acid substitutions at a single residue (A302) within the gamma-aminobutyric acid (GABA) receptor sub-unit named Rdl in the cat flea *Ctenocephalides felis* for 8 of the 9 laboratory strains that were tested (C. BASS *et al.* 2004). |
| Part of a coding region | Various amino acid substitutions in the DIII and DIV pore loops of the sodium channel Nav1.4 explain tetrodotoxin resistance in newts, snakes, pufferfishes (G. TOLEDO *et al.* 2016) and to red tide toxin-resistance in clams (V.M. BRICELJ *et al.* 2005). |
| Enhancer | Pelvic loss in sticklebacks originating from 13 different lakes is due to various deletions with distinct breakpoints of the same *pitx1* enhancer region in 9 cases and to other yet unknown genetic change(s) in 4 cases (Y.F. CHAN *et al.* 2010). |
| Gene | The gene *svb* was successfully predicted to cause an evolutionary loss of trichomes in *Drosophila ezoana* (FRANKEL, WANG and STERN 2012).<br>The gene *WntA* was successfully predicted to cause wing color pattern variation across many *Heliconius* species and populations (A. MARTIN *et al.* 2012), and beyond (J.R. GALLANT *et al.* 2014). |
| Gene Family, Gene Paralog | Interspecific changes in anthocyanin pigment intensity in flowers was successfully predicted to involve preferentially mutations in transcription factor genes of the MYB family (STREISFELD and RAUSHER 2011). |
| Genetic Pathway | It was successfully predicted that flower color evolution in *Penstemon barbatus* was caused by the inactivation of one of the candidate anthocyanin pathway genes, *F3'H*, *F3'5'H* and *DFR* (WESSINGER and RAUSHER 2014). |
| Gene Ontology | Animal morphological evolution is predicted to involve preferentially signaling ligand encoding genes, i.e. genes associated with the gene ontology (GO) terms “extracellular region” and “receptor binding” (MARTIN and ORGOGOZO 2017). |
| Broader Class of Mutation | Cis-regulatory mutations are the predominant mechanism underlying the evolution of morphology in animals (CARROLL 2008; STERN and ORGOGOZO 2008). |

## The causes of genetic repetition

In general, authors put forward several arguments for the over-representation of certain genetic loci for phenotypic traits of interests and suggest that their combination explains repeated evolution. Arguments for the importance of cis-regulatory mutations versus coding changes are not reviewed here and can be found in CARROLL 2000, CARROLL 2008, DAVIDSON 2006, Wray 2007, STERN and ORGOGOZO 2008, STERN and ORGOGOZO 2009, B.-Y. LIAO, M.-P. WENG and J. ZHANG 2010, STERN 2011.

Several non-exclusive hypotheses have been proposed to explain why some genetic changes repeatedly drive certain phenotypic changes or adaptations during evolution (Table 2). A first category of explanation can be attributed to mutational bias: certain mutations are more likely to occur than others. For instance, a region that is prone to structural variation or transposon insertion is likely to undergo repeated rearrangements, thus facilitating certain gene-to-phenotype changes (CHAN *et al.* 2010). Adaptation to high-altitude in Andean house wrens and hummingbirds has been associated with single point mutations in the *βA-globin* gene and these mutations appear to lie within CpG sites, which are known to be chemically instable and highly mutable (S.C. GALEN *et al.* 2015; A. STOLTZFUS and D.M. MCCANDLISH 2015; M. LYNCH and B. WALSH 2007). A recent study of various high-altitude species (C. NATARAJAN *et al.* 2016) discovered other mutations in *βA-globin* that increase affinity for oxygen, indicating that other genetic paths are theoretically possible. In Andean house wrens the spontaneous mutation rate appears to have biased evolution towards certain genetic paths. Spontaneous mutation rates are higher for transitions (A↔G or C↔T) than for transversions (M. LYNCH and B. WALSH 2007). Using this property as a test case for investigating the role of mutation biases, Stoltzfus and McCandlish compiled from existing literature a list of putatively adaptive amino-acid changes that have evolved in parallel in natural or experimental contexts. They found a fourfold excess of transitions over transversions (A. STOLTZFUS and D.M. MCCANDLISH 2017), suggesting that the repeatability of adaptive coding changes is at least partly explained by biases in the mutations. This is consistent with a “first come, first served” model where even if a number of possible mutational paths to adaptation exist, the ones that are more likely to emerge in the first place are more accessible to selection, and are thus repeatedly observed when the environmental challenge is replicated.

**Table 2.**
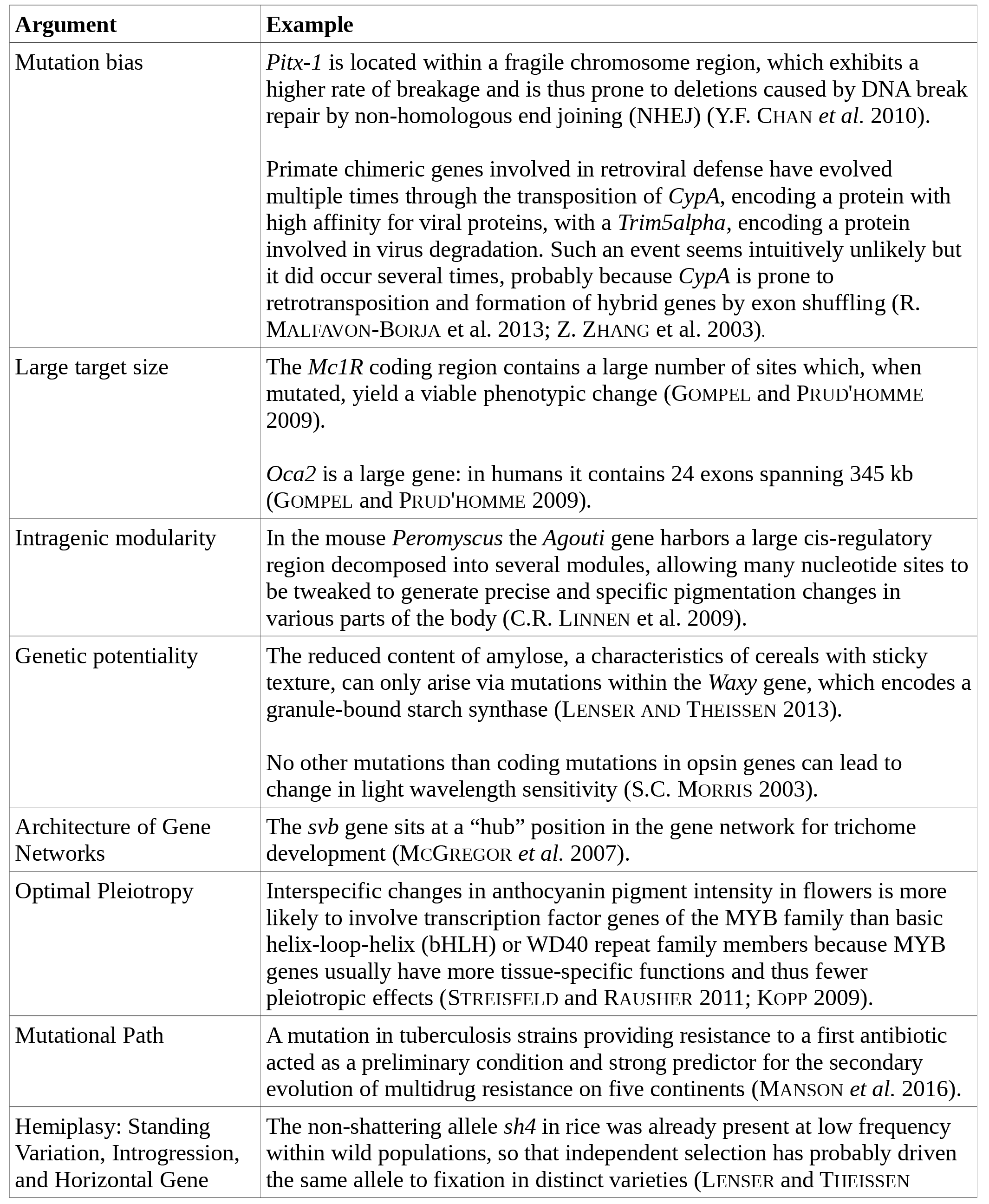
**Non-exhaustive list of arguments that have been proposed to explain hotspot genes.**

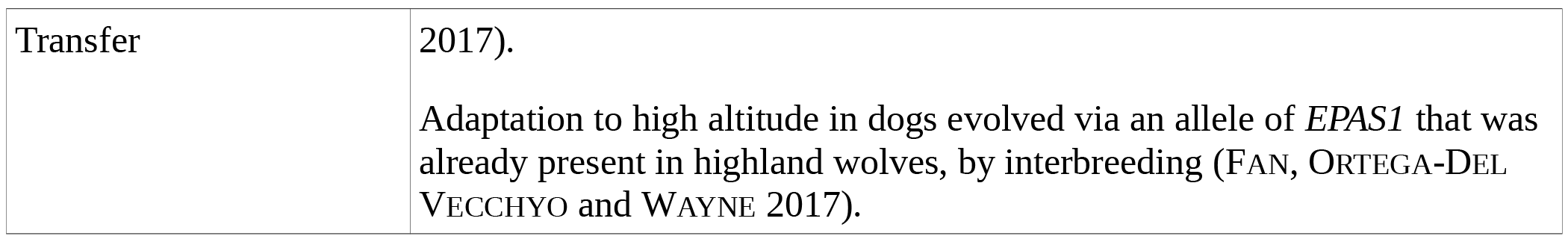

Two other categories of explanations (named “Intragenic Modularity” and “Large Target Size” in Table 2) are related to the fact that compared to other loci that may provide mutational paths to the considered phenotypic change, a given gene may be favored because of its properties at the DNA sequence level: a large intergenic region that can provide relatively more possibilities for a modular control, or a large coding region allowing many amino-acid sites to be tweaked.

A third category of explanation deals with gene function itself rather than just the physical properties of the stretch of DNA hosting their information. Kopp used the term “Optimal Pleiotropy” to propose that only certain genes may host the potential for tuning a given phenotype without deleterious effects. In this concept, the emphasis is on the capacity of certain genes to yield specific effects, whether they simply have a limited number of roles, or on the contrary, a large number of roles but sufficient modularity to allow genetic uncoupling of these roles (A. KOPP 2009). Similarly, the “Architecture of Gene Networks” may highlight “hub” or “input/output” genes that are more likely to coordinate a cascade of changes and thus, drive effects accessible to selection (MCGREGOR *et al.* 2007; STERN and ORGOGOZO 2008).

A fourth mechanism deals with “Permissive Mutations” that may relax the constraints preventing a given change and thus open a new valley in the adaptive landscape. This phenomenon was reviewed recently (J.F. STORZ 2016) and has been mostly studied in the framework of protein evolution (R.D. TARVIN *et al.* 2017; J.D. BLOOM, L.I. GONG and D. BALTIMORE 2010; A. MANSON *et al.* 2017; T.N. STARR, L.K. PICTON and J.W. THORNTON 2015). The predictability is of the form “if change A has happened, and then change B is likely as well because we observed that A is often followed by B”. In principle, such phenomenon could also apply to genome-wide epistasis, with mutational paths being contingent on allelic states present at distant loci on the genome, as has been observed in “Evolve and Resequence” experiments (J.B. ANDERSON *et al.* 2010; A. LONG *et al.* 2015).

Finally, another explanation underlines the higher capacity of certain derived alleles to circulate among the branches of a species phylogeny, due to various mechanisms such as incomplete lineage sorting or standing genetic variation (*eg.* C.T. MILLER *et al.* 2007), genetic introgressions in intermixing populations or closely related species (*eg.* J. ENCISO-ROMERO *et al.* 2017; Z. FAN, D. ORTEGA-DEL VECCHYO and R.K. WAYNE 2017; L.T. DUNNING *et al.* 2017), or even horizontal gene transfers between distant branches of the tree of life (*eg*. F.W. Li *et al.* 2014; J.A. METCALF *et al.* 2014; J. ROPARS *et al.* 2015). These cases, named “collateral evolution” or “hemiplasies” (reviewed in STERN 2013; MARTIN and ORGOGOZO 2013) contrast with *stricto sensu* occurences of genetic parallelism (R.W. SCOTLAND 2011; STERN 2013; STORZ 2016) because the alleles are identical-by-descent rather than identical-by-state. The derived phenotypes may be “convergent”, *ie.* show a pattern of homoplasy due to a discontinuity on a phylogeny, but the genotypes are not, hence representing a so-called “hemiplasy” (J.C. AVISE and T.J. ROBINSON 2008). As our understanding of gene flow is rapidly improving in the phylogenomic era, it is likely that the *a priori* observation of connectivity between two lineages will increase our capacity to make predictions on causal mechanisms of gene-to-phenotype change (R.W. WALLBANK *et al.* 2016).

In summary, predictable genetic changes may be *1)* under the influence of a mutational bias ( site-specific rates of mutation); *2)* in a large locus, more prone to change due to a sequence size parameter (narrow mutational target size, independently of genetic function – large target size, intragenic modularity); *3)* in a gene that is inherently poised to tweak a given trait due to properties of its molecular function or regulatory interactions (narrow mutational target size, due to the structure of the genotype-phenotype map - Genetic potentiality, Architecture of Gene Networks, Optimal Pleiotropy); *4)* contingent upon the pre-existence of other changes in the genome (Mutational Path); 5) a by-product of allele sorting or transfer processes (Hemiplasy).

## Prediction accuracy

Good predictions are not only the ones with high inference gain, but also the ones with high *accuracy*. As a matter of fact, the prediction that tomorrow will be a sunny day is better with 95% accuracy than with 50% accuracy. Accuracy corresponds to the ratio of correct predictions to the total number of cases evaluated. How accurate are the predictions about the genetic loci of evolution? Some appear to be 100% accurate whereas others are not. Pigmentation evolution in flies was predicted to involve mostly cis-regulatory mutations (CARROLL 2008). A recent review compiles 32 cases of pigmentation evolution in various *Drosophila* species: all of them are caused by cis-regulatory mutations and none are caused by coding changes (J.H. MASSEY and P.J. WITTKOPP 2016). If we suppose that existing approaches are not biased towards cis-regulatory changes, then it means that so far the prediction for cis-regulatory evolution is 100% accurate. Resistance to trimethoprim in *Escherichia coli* was correctly predicted to be associated with genes encoding dihydrofolate reductase enzymes in 308 of the 320 tested resistant strains, thus giving an accuracy of 96% (A. BROLUND *et al.* 2010). An examination of 192 worldwide populations of *Arabidopsis thaliana* exhibiting natural variation in flowering time found that approximately 70% of the early-flowering strains carried deleterious mutations in the hotspot gene *FRIGIDA* (C. SHINDO *et al.* 2005). The prediction that variation in flowering time in *A. thaliana* should involve deleterious mutations in *FRIGIDA* has thus an estimated accuracy of 70%. In tetrapods, the *MC1R* receptor and its antagonist *Agouti* together account for 54% of the 206 published cases of pigmentation variation (MARTIN and ORGOGOZO 2017). This percentage is inflated by a “caveman effect”, which is a type of sampling bias: these two genes might be called “pigmentation hotspot genes” because there are precisely the loci researchers look at first when digging for genetic changes driving pigment variation. Nonetheless, the 54% value gives a maximum estimate of the accuracy of the prediction for a commonly studied trait.

The prediction that genes encoding signaling ligands should be responsible for morphological evolution is one of the less accurate predictions that have been presented. According to experimental data, about 20% of the cases where an animal morphological difference has been mapped to a gene involve a signaling ligand gene (80/391, MARTIN and ORGOGOZO 2017, https://www.gephebase.org). In sticklebacks, 14 genome-wide QTL studies ended with the identification of the causal gene and 4 of them identified a secreted ligand gene, thus giving an estimated accuracy of 28%. The ligand prediction is still better that the null model of each gene having an equal probability of being responsible for the phenotypic change, as ligand genes represent less than 5% of the total number of genes in a genome (MARTIN and ORGOGOZO 2017). A more accurate formulation of the ligand prediction is thus that for animal morphological evolution signaling ligand genes are over-represented compared to their proportion in genomes.

Interestingly, the accuracy of actual predictions does not appear to be determined by the strength of the arguments that have been proposed to substantiate them. A wealth of arguments have been proposed for the importance of cis-regulatory mutations in morphological evolution (CARROLL 2000; PRUD'HOMME, GOMPEL and CARROLL 2007; WRAY 2007) and yet this prediction does not appear to be accurate for plant morphological evolution (STREISFELD and RAUSHER 2011).

## Inference gain varies with phenotypes

Certain phenotypic traits are associated with predictions of high inference gain while other phenotypic changes call for low inference gain predictions. Predictions about metabolic activity or resistance to particular molecules appear to have more inference gain than predictions about morphological differences. For example, evolution of C4 photosynthesis can be associated with a few specific amino acid changes in the *PEPC* gene (CHRISTIN *et al.* 2007) and antifolate resistance in *Plasmodium falciparum* with only 6 mutations in the dihydrofolate reductase (*DHFR*) locus (M.S. COSTANZO and D.L. HARTL D.L. 2011). In contrast, pigmentation pattern evolution can be caused by mutations in at least 10 genes in *Drosophila* flies and 13 genes in Vertebrates (Table 3). Importantly, additional data about the phenotypic change of interest can help narrow down the number of candidate genetic loci. Coat-darkening phenotypes in natural populations of Vertebrates have been associated to only two of these pigmentation genes, the *Agouti* signaling protein (*Agouti*) and melanocortin-1 receptor (*Mc1r*). So these are the best candidates genes for natural coat-darkening phenotypes. Of note, spontaneous coat-darkening phenotypes in mice have also been associated with mutations in two other genes, *attractin* (*Atrn*), and *mahogunin* (*Mgrn*) (E.P. KINGSLEY *et al.* 2009). Moreover, knowing whether the coat-darkening phenotype is dominant or recessive reduces the number of candidate mutations further: a gain-of-function in *Mc1R* is inferred for dominant phenotypes and a loss-of-function in *Agouti* for recessive traits (E. EIZIRIK *et al.* 2003). The better characterized the phenotypic change, the more inference gain one can have. For example, pelvic-reduced sticklebacks were found to exhibit a left-right directional asymmetry of pelvic bones, as in P*itx1-*null mice, thus strengthening the prediction that the underlying gene should be *Pitx1* (M.D. SHAPIRO *et al.* 2004, Y.F. CHAN *et al.* 2010).

**Table 3.**
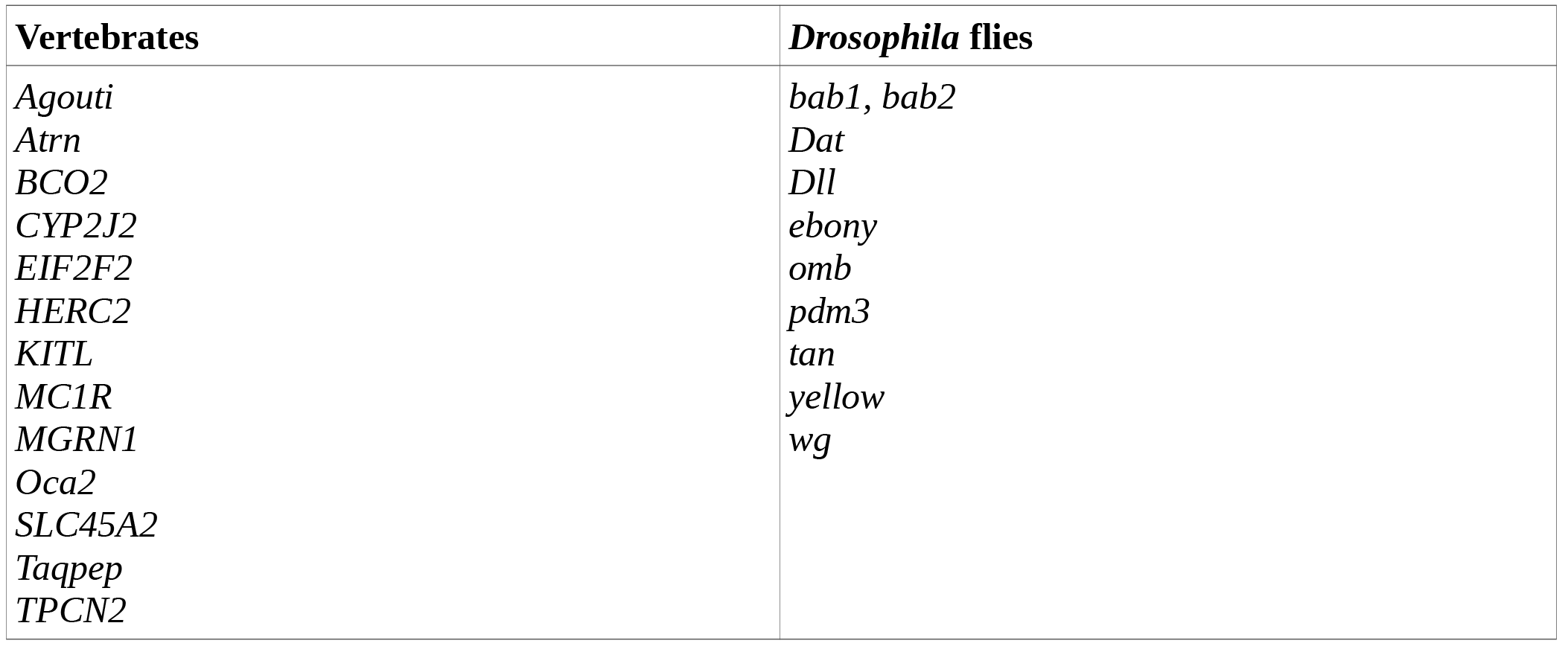
**Non-exhaustive list of genes responsible for evolution of pigmentation pattern in natural populations in Vertebrates and *Drosophila flies***. See https://www.gephebase.org for details.

Rockman and others have argued in favor of a polygenic model behind complex traits (R.A. Fisher 1918; M.V. ROCKMANN 2012; E.A. BOYLE, Y.I. LI and J.K. PRITCHARD 2017), suggesting that the genetic loci underlying certain complex traits may not be predictable. We do not doubt the generality of this view, but it is useful to stress that traits assumed to be multigenic are sometimes found to be oligogenic (S. MAKVANDI-NEJAD *et al.* 2012). Focusing on large-effect loci can facilitate the discovery and identification of some of the genetic causes of phenotypic variation. This said, it remains critical to look for alternative mutations and avoid the pitfall of ascertainment bias. The power of contemporary association mapping in naturally variable populations and of genome-wide genotyping for QTL analysis shall facilitate our escape from the low-hanging large-effect loci and help us draw a balanced view of genetic predictability in the next decades.

## Does prediction accuracy vary with time scale and taxonomic range?

As the taxon of interest becomes more distant from the set of species from which genetic knowledge is available and from which predictions are elaborated, predictions may be less likely. For example, table 3 shows that genes involved in pigmentation evolution in Vertebrates cannot be used as candidate genes for pigmentation evolution in flies. C4 photosynthesis evolved many times independently in grasses and sedges through mutations in phosphoenolpyruvate carboxylase (PEPC) via a limited number of amino acid positions and the distribution of sites that have repeatedly mutated differ significantly between grasses and sedges, indicating that the genetic basis of C4 photosynthesis evolution is slightly different between taxa (CHRISTIN *et al.* 2007). A meta-analysis of ~25 cases (CONTE *et al.* 2012) suggested that probability of gene reuse declines with divergence time between the two taxa under consideration. However, this trend was not reproduced with a larger dataset (118 cases, Fig. 6 in Gallant *et al.* 2014). Therefore, we cannot conclude from current data that independent evolution of the same phenotype is more likely to involve mutations in the same genetic locus when taxa are closely related than when they are distantly related.

## Conclusion

Even though predictions about the loci of past evolution do not rely on advanced theoretical models, they have proved relatively accurate so far. Predicting the mutations of the past can help not only to understand the mechanisms of evolution, but also to genetically-engineer domesticated species and to infer the mutations that will occur in pathogenic microorganisms.

## Acknowledgements

We thank S. Caianiello, S. Campanella, D. Ceccarelli, G. Frezza and E. Gagliasso for organizing the 2017 conference and for encouraging the writing of this review. We also thank A. Stoltzfus, G. Conte B. Morizot and A. Love for discussions, and the Courtier-Orgogozo team and the labex “Who am I?” for providing a stimulating environment. The research leading to this paper has received funding from the European Research Council under the European Community's Seventh Framework Program to VCO (FP7/2007-2013 Grant Agreement #337579) and from the John Templeton Foundation to AM and VCO (JTF award #43903).

